# Genome sizes of animal RNA viruses reflect phylogenetic constraints

**DOI:** 10.1101/2024.11.01.621630

**Authors:** Kosuke Takada, Edward C. Holmes

## Abstract

Animal genomes are characterized by extensive variation in size. RNA viruses similarly exhibit substantial genomic diversity, with genome lengths ranging from 1.7 to 64 Kb. Despite the myriad of novel viruses discovered by metagenomics, we know little of the factors that shape the evolution of the genome size in RNA viruses. We analyzed the variation in genome sizes across orders and families of animal RNA viruses. We found that RNA viruses can have highly variable genome sizes within and among orders, with the *Nidovirales* (including the *Coronaviridae*) having both significantly larger genomes and a greater range of genome sizes than other orders. In the *Bunyavirales, Amarillovirales, Nidovirales* and *Picornavirales* the genome sizes of invertebrate-associated RNA viruses were significantly larger than those that infect vertebrates, in contrast to their animal hosts in which vertebrates commonly have larger genomes than invertebrates. However, in the *Mononegavirales*, vertebrate viruses were significantly larger than those viruses associated with invertebrates. There were similarly complex associations between genome size and patterns of genome segmentation. In the *Bunyavirales, Reovirales*, and *Nidovirales*, viruses with segmented genomes or that possessed a large number of segments, had significantly larger genome sizes that viruses with non-segmented genomes or a small number of segments, while in the *Articulavirales* there were no significant differences in genome size among viruses possessing any number of genome segments. More broadly, our analysis revealed that taxonomic position (i.e., RNA virus order) had a greater impact on genome size than whether viruses infected vertebrates or invertebrates or their pattern of genome segmentation. Hence, the phylogenetic constraints on genome size are of sufficient magnitude to shape other aspects of virus evolution.

## Introduction

Animal genomes are characterized by extensive variation in size. In general, vertebrate genomes are larger and more complex than those of invertebrates (Gregory 2005), varying from around 400 million base pairs (Mb) in some fish (e.g., *Takifugu rubripes*) (Aparicio et al. 2002; Gregory 2024) to billions base pairs (Gb) in mammals and birds (Gregory 2005; Venkatesh et al. 2000). The largest vertebrate genome reported to date is ∼130 Gb in the lungfish *Protopterus aethiopicus*, while genomes of ∼117 Gb are found in the amphibians *Necturus lewisi* and *Necturus punctatus*. Repetitive DNA sequences such as transposable elements (i.e., DNA transposons and retrotransposons) are a major contributor to the large genome sizes of some vertebrate species, and associated with expansions in genome size and complexity (Biscotti et al. 2015; Canapa et al. 2015; Kidwell 2002). In contrast, invertebrate genome sizes may be only around 20 Mb in the nematode *Pratylenchus coffeae*, ∼180 Mb for the arthropod *Drosophila melanogaster*, but over 1 Gb for certain insects and marine invertebrates (Canapa et al. 2015; Gregory 2024). It is important to note, however, that proportionally far fewer genomes have been sequenced from invertebrates than vertebrates. Despite these sampling biases, it is clear that there are marked differences in the size of animal genomes.

Animals (*Metazoa*) evolved from a single-celled ancestor more than 635 million years ago, an evolutionary transition that was associated with a dramatic increase in phenotypic diversity, including the rapid diversification of multiple animal phyla (Davidson and Erwin 2009; dos Reis et al. 2015; Ros-Rocher et al. 2021). A key evolutionary innovation associated with the later rise of the vertebrates was the adaptive immune system, first appearing in jawed fish about 500 million years ago, and leading to major differences in how vertebrates respond to viral infections compared to their invertebrate ancestors (Flajnik and Kasahara 2010). In particular, while the invertebrate immune response is innate and usually stereotypic, that of vertebrates is more complex and has a strong adaptive capacity, allowing for a more rapid and targeted immune response when pathogens are repeatedly encountered (Boehm 2012; Du Pasquier 2001). It might also be expected that such a major transition in host evolution, concordant with the evolution of the vertebrates, would impact the evolution of the viruses that infect them, including the size and complexity of their genomes. For example, it is has been hypothesized that the evolution of the adaptive immune system would select for smaller virus genomes as this would theoretically result in fewer immune targets (Harvey and Holmes 2022).

Viruses are similarly characterized by considerable genomic diversity. Genome sizes in RNA viruses range from only around 1.7 Kb (Hepatitis delta virus, family *Kolmioviridae*) to more than 64 Kb (members of the order *Nidovirales*) (Gorbalenya et al. 2006; Neuman et al. 2024). Although metagenomic sequencing of diverse animal taxa in recent years has led to a gradual increase in our knowledge of genome sizes in RNA viruses, particularly in the *Nidovirales* (Neuman et al. 2024), these are usually still far smaller than those of DNA viruses. Indeed, the genome sizes of DNA viruses range from ∼1.8 Kb for the (single-stranded) circoviruses to ∼2.5 Mb for the pandoraviruses (Philippe et al. 2013).

The constraint on the maximum length of RNA virus genomes has commonly been associated with the low fidelity of their replication process, central to which is a RNA-dependent RNA polymerase (RdRp) that lacks proofreading ability and hence results in a high mutation rate per genome replication (Holmes 2009). Indeed, there is a broad inverse correlation between genome size and mutation rate (Duffy et al. 2008; Sanjuan et al. 2010), such that an increase in virus genome size requires a reduction in mutation rate per nucleotide to limit the accumulation of deleterious mutations that result in major fitness losses (Holmes 2003). An exception that goes some way to proving this rule are the *Coronaviridae*, again members of order *Nidovirales*, that possess a 3′-to-5 exoribonuclease (ExoN) protein (i.e., N-terminal domain of nonstructural protein 14) that provides a form of RNA proof-reading that reduces error rates and hence facilitates the evolution of longer genomes (Eckerle et al. 2010; Gorbalenya et al. 2006; Liao et al. 2023). Although it is unclear whether similar error-correction mechanisms exist in other RNA viruses with large genome sizes, it is noteworthy that a 39.8 Kb virus was recently identified within the *Flaviviridae* (order *Amarillovirales*) that does not encode an exoribonuclease or known error-correction protein (Petrone et al. 2024).

It may also be that genome sizes of RNA viruses differ according to their type of genome organization. In particular, it has been suggested that segmented RNA viruses may have larger genomes than non-segmented (i.e., single segment) RNA viruses because this reduces mutation accumulation by facilitating the more efficient selective removal of deleterious mutations through genomic reassortment (Chao 1990; Pressing and Reanney 1984). Indeed, one study revealed that the average size of segmented RNA virus genomes was slightly larger, at ∼11 Kb, than that of non-segmented RNA viruses (9 Kb, although excluding members of the *Coronaviridae* that possess ExoN domains) (Holmes 2009). Currently, the largest reported segmented RNA viruses are some bi-segmented members of the order *Nidovirales* with genomes that exceed 36 Kb (Neuman et al. 2024). To date, however, there has been no systematic assessment of the relationship between the patterns of genome segmentation and genome size.

Despite the huge expansion in the known virosphere that has resulted from the development of metagenomic sequencing, we currently have little understanding of the factors that shape the evolution of the genome size of RNA viruses, particularly if they differ fundamentally between hosts – vertebrates and invertebrates – that themselves differ in genome size and complexity. To better understand the factors responsible for the range of genome sizes of RNA viruses, we performed a detailed analysis of the variation in genome sizes across different orders and families of RNA viruses, and examined the differences between non-segmented and segmented viruses and those that infect vertebrates or invertebrates.

## Results

The International Committee on Taxonomy of Viruses (ICTV) currently approves eight primary taxonomic ranks for RNA viruses (realm, kingdom, phylum, class, order, family, genus, and species in descending order) and seven secondary ranks (subrealm, subkingdom, subphylum, subclass, suborder, subfamily, and subgenus). The realm *Riboviria* includes the kingdom *Orthornavirae*, that comprise RNA viruses encoding an RNA-dependent RNA polymerase (RdRp), and the kingdom *Pararnavirae* that consists of reverse-transcribing viruses that encode a reverse transcriptase. The *Orthornavirae* currently includes four phyla – *Lenarviricota, Pisuviricota, Kitrinoviricota, Duplornaviricota* and *Negarnaviricota* – that contain animal RNA viruses, and the phylum *Lenarviricota* that likely comprise bacterial, fungal and protist RNA viruses.

To reveal the diversity of genome lengths at the level of virus order in the diverse animal-derived RNA viruses found within each of the four *Riboviria* phyla (i.e., *Pisuviricota, Kitrinoviricota, Duplornaviricota* and *Negarnaviricota*), excluding those associated with plants and other hosts, we compared the size of complete genomes of viruses registered in the Virus Genome Resource (https://www.ncbi.nlm.nih.gov/genomes/GenomesGroup.cgi?taxid=10239&sort=taxonomy). This analysis revealed considerable variation in viral genome sizes between orders (Fig. 1A, Table 1). In the 18 viral orders analyzed here, the *Durnavirales* had the smallest genome sizes (mean of 4,148 ± 268.1 nt), while the *Nidovirales* had the largest (mean of 26,178 ± 7,457 nt, but up to 64,336 nt) (Table 1). Indeed, the median and both lower and upper 95% confidence intervals of the mean were the higher in the *Nidovirales* than other orders (median of 27,464 nt, lower 95% confidence interval =25,002 nt, upper 95% confidence interval = 27,354 nt) (Table 1). The orders *Reovirales* and *Amarillovirales* also contained viruses with large genome sizes (up to 29,174 nt and 39,834 nt, respectively), with the *Amarillovirales* characterized by a highly skewed distribution toward smaller genome sizes and with extreme values at the distribution tails (range of 31,266 nt; from 8,568 nt to 39,834 nt, skewness of 4.094, kurtosis of 21.67) (Table 1). The *Nidovirales* similarly have a very wide range of genome sizes, spanning 51,632 nt; from 12,704 nt in Equine arteritis virus to 64,336 nt in Crassostrea gigas nidovirus (Fig. 1A, Table 1, Supplementary Table 1).

**Table 1.**
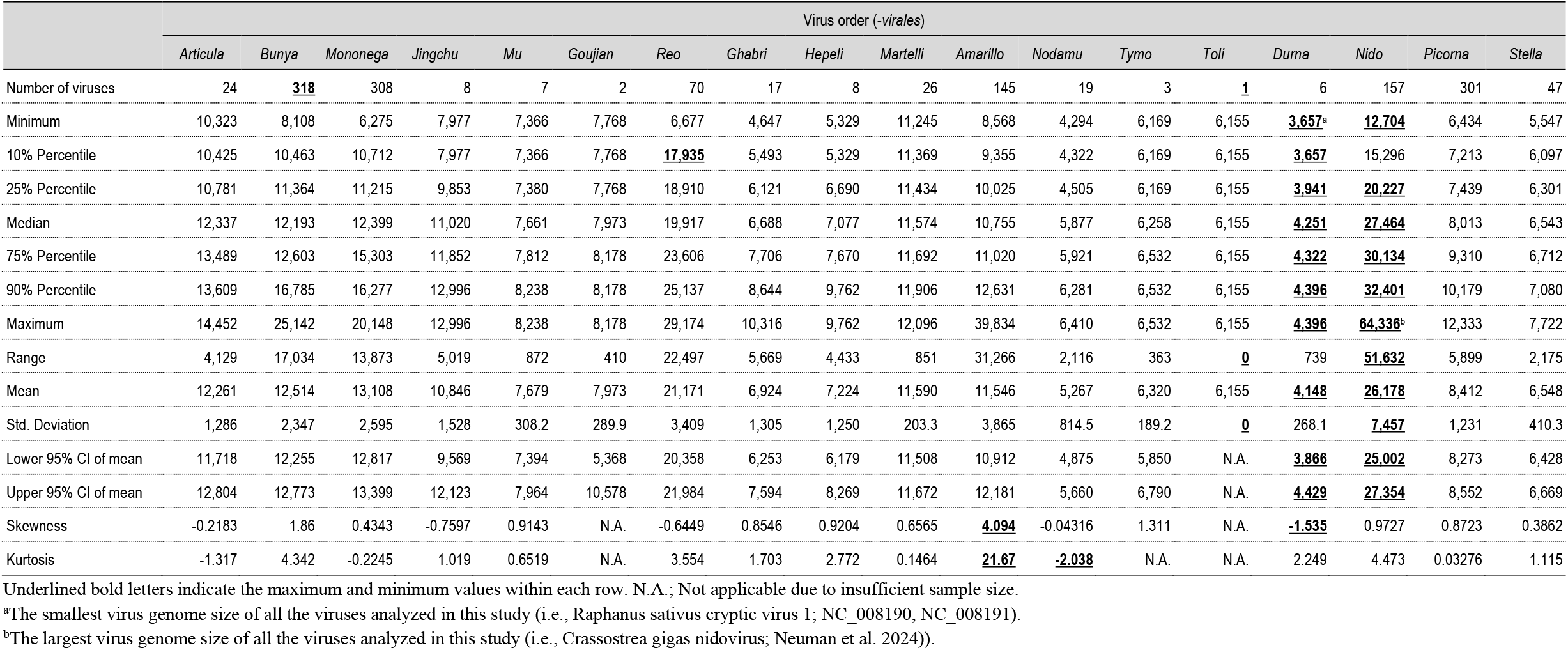
Comparison of genome sizes of animal viruses within orders of RNA viruses.

**Figure 1.**
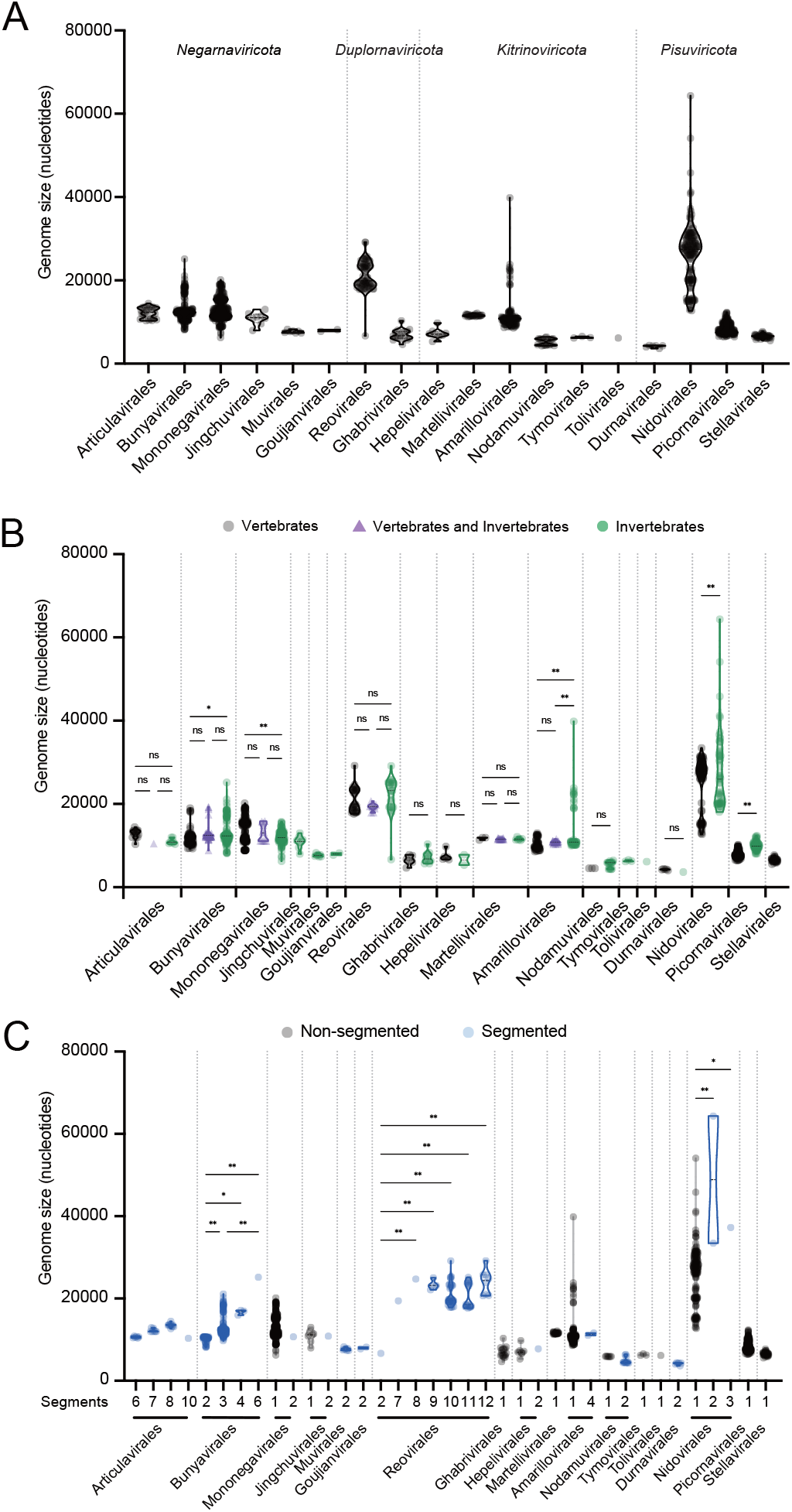
Genome sizes of animal RNA viruses within each viral order. (A) Genome sizes of animal RNA viruses belonging to each order studied are shown in the truncated violin plot. (B) The genomes size of vertebrate- and invertebrate-associated RNA viruses in each order are shown in the truncated violin plot. Vertebrate-associated RNA viruses, RNA viruses associated with both vertebrates and invertebrates, and invertebrate-associated RNA viruses are shown in black, purple and green, respectively. (C) The genome sizes of non-segmented and segmented RNA viruses belonging to each viral order are shown in the truncated violin plot. Non-segmented and segmented RNA viruses are shown in black and blue, respectively. The number of genome segments is indicated above the name of each viral order (in which “1” denotes a non-segmented genome). In all cases, the mean values within each viral order were compared by using a one-way ANOVA, followed by Šídák’s multiple comparisons test (ns = not significant with *P* >0.05; \**P* < 0.05; \*\**P* < 0.01).

To determine whether there were differences in the variation in genome size between viral orders, we performed the mean-based Levene’s test of equality of variance (Draper and Hunter 1969), excluding *Goujianvirales* and *Tolivirales* which only contained two and one viruses, respectively (Table 1). The distance (i.e., absolute deviation) from the mean of each virus genome was calculated for each virus in each order, which were then compared between orders. As expected, the *Nidovirales* had significantly larger mean genome sizes than other orders (*p* < 0.01) (Supplementary fig. 1). In addition, the mean genome size of the *Mononegavirales* was significantly larger than that of the *Bunyavirales, Martellivirales*, and *Stellavirales* (*p* = 0.0184, *p* = 0.003, *p* <0.0001 and *p* <0.0001, respectively), while that of the *Reovirales* was significantly larger than the *Bunyavirales, Martellivirales, Picornavirales* and *Stellavirales* (*p* = 0.0016, *p* = 0.0001, *p* <0.0001 and *p* <0.0001, respectively) (Supplementary fig. 1). There were also significant differences in mean genome sizes between the *Amarillovirales* and *Martellivirales, Amarillovirales* and *Picornavirales*, and *Amarillovirales* and *Stellavirales* (*p* = 0.0124, *p* = 0.0016, *p* = 0.0008, respectively) (Supplementary fig. 1). Hence, these results indicate that the genome sizes of animal-derived RNA viruses vary widely both within and between viral orders.

Since characteristics of viruses are strongly dependent on their host and how they respond to viral infections, it is conceivable that viral genome size could be affected by the major host group they infect. We therefore analyzed whether the animal RNA viruses analyzed here were associated with either vertebrates or invertebrates. Accordingly, some viral orders (*Articulavirales, Bunyavirales, Mononegavirales, Reovirales, Ghabrivirales, Hepelivirales, Martellivirales, Amarillovirales, Nodamuvirales, Durnavirales, Nidovirales* and *Picornavirales*) contained both vertebrate- and invertebrate-associated RNA viruses (Fig. 2). The same was true at the family level, where the *Orthomyxoviridae, Nairoviridae, Peribunyaviridae, Phenuiviridae, Rhabdoviridae, Sedoreoviridae, Spinareoviridae, Totiviridae, Togaviridae, Flaviviridae, Nodaviridae* and *Tornidovirineae* contained both vertebrate- and invertebrate-associated RNA viruses (Fig. 2).

**Figure 2.**
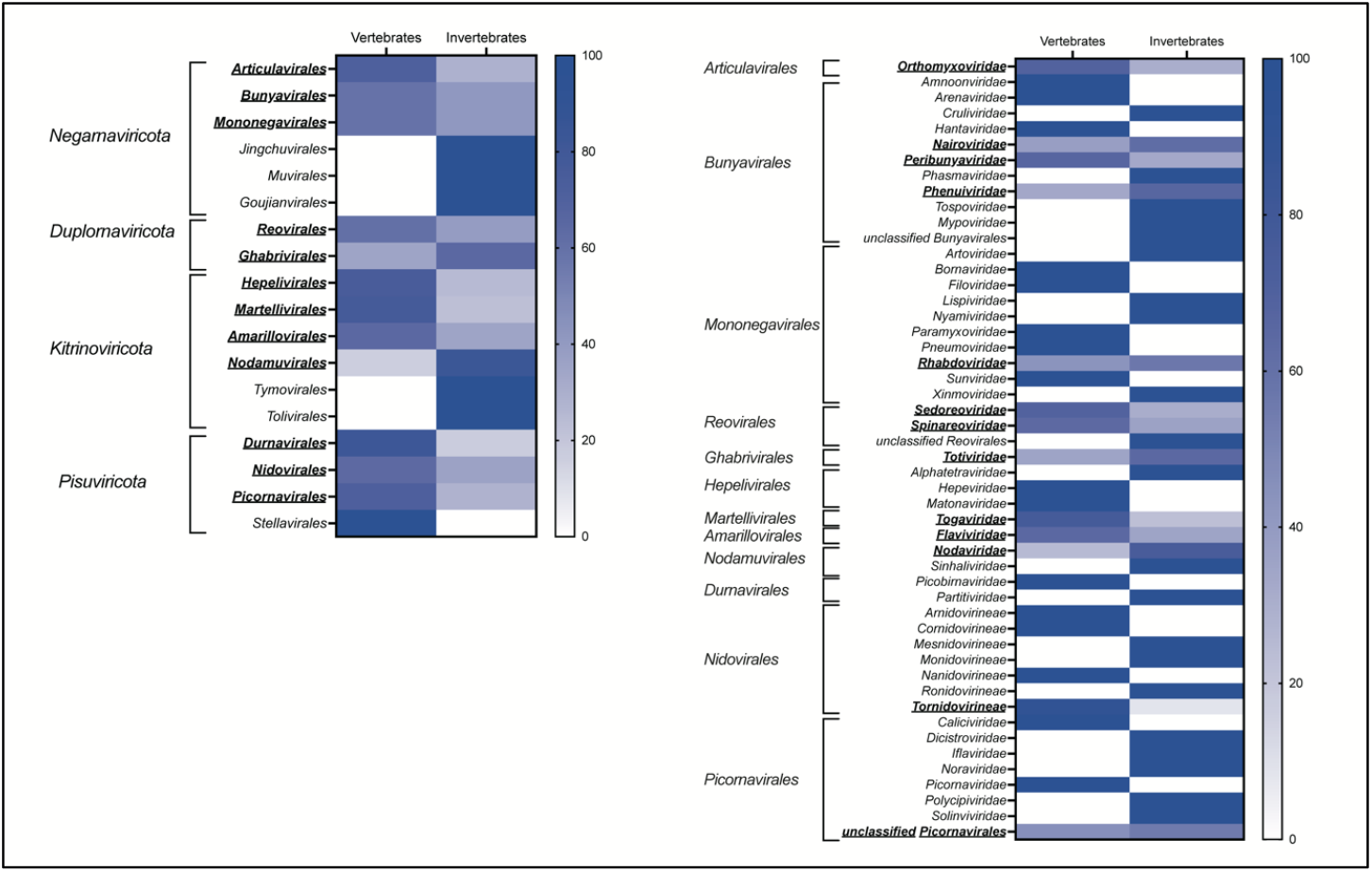
Vertebrate and invertebrate-associated RNA viruses in each viral order and viral family. (A) Heatmap showing the proportion of vertebrate- or invertebrate-associated RNA viruses in each viral order. Underlined bold letters indicate the virus orders that include both vertebrate- and invertebrate-associated viruses. (B) Heatmap showing the proportion of vertebrate- or invertebrate-associated RNA viruses in each virus family in the viral orders that contained both types of virus. Underlined bold letters indicate the virus families that include both vertebrate- and invertebrate-associated viruses.

We similarly compared the genome sizes of the vertebrate- and invertebrate-associated RNA viruses within each virus order. Notably, in the *Bunyavirales, Amarillovirales, Nidovirales* and *Picornavirales*, invertebrate-associated viruses had significantly larger genome sizes than vertebrate viruses (*p* = 0.0135, *p* < 0.0001, *p* < 0.0001 and *p* < 0.0001, respectively) (Fig. 1B). Conversely, in the *Mononegavirales*, the genome size of vertebrate-associated viruses was significantly larger than that of invertebrate viruses (*p* < 0.0001) (Fig. 1B). This was the only group in which vertebrate viruses were longer than those in invertebrates. Non-significant differences between vertebrate and invertebrate viruses were observed in the remaining inter-order comparisons. Similarly, we compared the genome sizes of vertebrate- and invertebrate-associated viruses within each virus family in those orders that showed differences in genome size between these major host groups. All viruses in the *Amarillovirales* analyzed here belonged to *Flaviviridae* (i.e., unclassified *Amarillovirales* were excluded from our analysis), within which the invertebrate-associated viruses had significantly larger genome sizes than those associated with vertebrates (*p* < 0.0001) (Supplementary fig. 2). In the *Nidovirales*, the *Arnidovirineae*, which are composed solely of vertebrate-associated viruses, had smaller genomes than other families in that order that are composed exclusively of invertebrate-associated viruses, while in the *Tornidovirineae* there were no significant differences in the size of genome between vertebrate- and invertebrate-associated viruses (*p* = 0.709) (Supplementary fig. 2). In the *Picornavirales*, the *Caliciviridae* and *Picornaviridae*, which are composed exclusively of vertebrate-associated viruses, had smaller genomes than other families in that order that are composed exclusively of invertebrate-associated viruses (Supplementary fig. 2). In the *Phenuiviridae* (*Bunyavirales*) and *Rhabdoviridae* (*Mononegavirales*) there were no significant differences in the genome sizes of vertebrate- and invertebrate-associated viruses (*p* = 0.9186 and *p* = 0.0627, respectively) (Fig. 3A and 3B). Finally, in the *Mononegavirales*, the *Filoviridae, Paramyxoviridae, Pneumoviridae* and *Sunviridae*, which are exclusively comprised of vertebrate viruses, had larger genomes than other families in that order that only contained invertebrate viruses (Fig. 3B).

**Figure 3.**
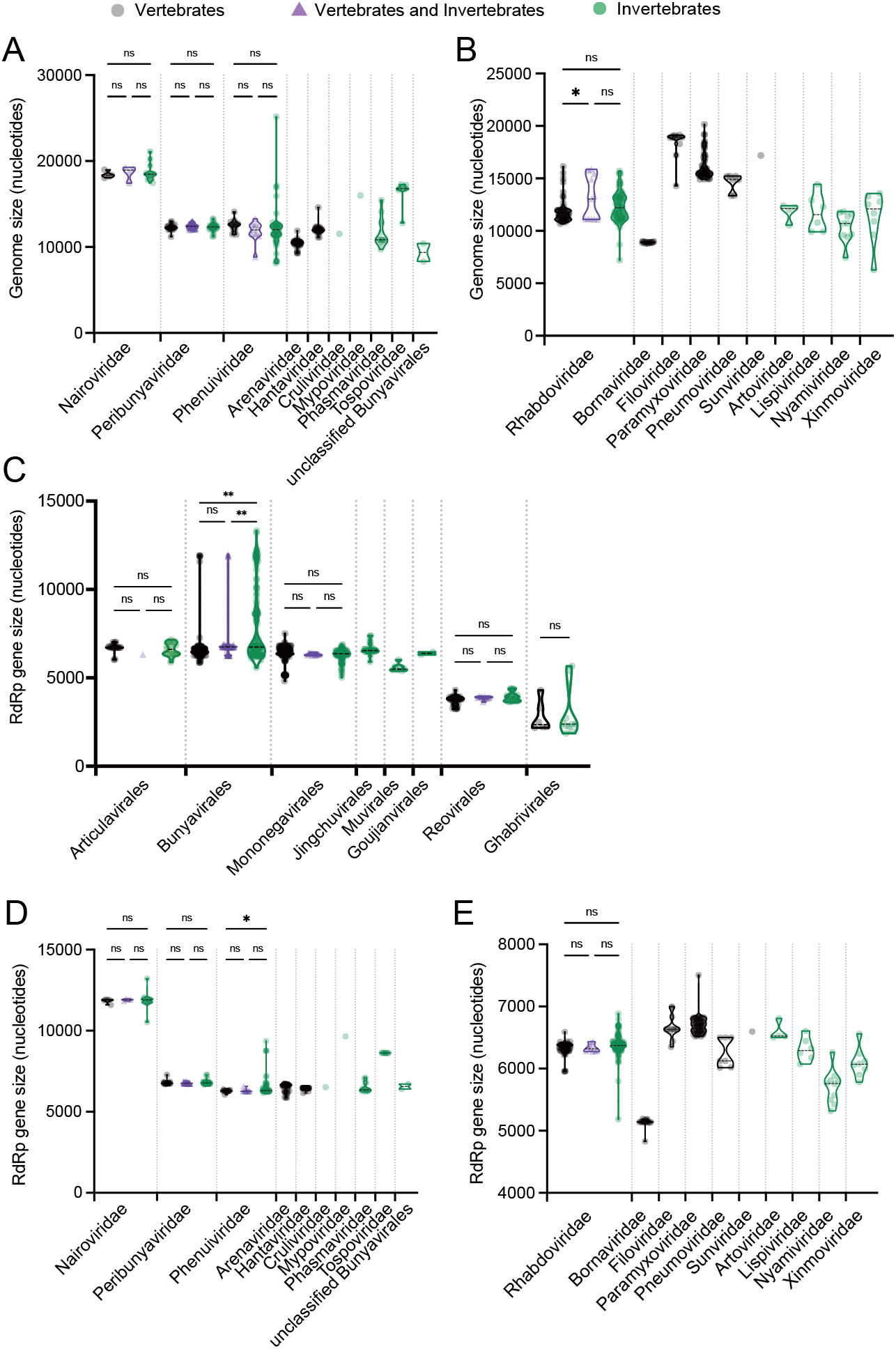
Genome sizes and RdRp gene lengths in animal RNA viruses. (A-B) Size of the genome of RNA viruses in each viral family belonging to the (A) *Bunyavirales* and (B) *Mononegavirales*. Vertebrate-associated RNA viruses, both vertebrate- and invertebrate-associated RNA viruses, and invertebrate-associated RNA viruses are shown in black, purple and green, respectively. (C) Size of the RdRp gene in each viral order within the *Negarnaviricota* and *Duplornaviricota*. Vertebrate-associated RNA viruses, RNA viruses associated with both vertebrates and invertebrates, and invertebrate-associated RNA viruses are shown in black, purple and green, respectively. (D-E) Size of the RdRp gene in each viral family within the *Bunyavirales* (D) and *Mononegavirales* (E). Vertebrate-associated RNA viruses, both vertebrate- and invertebrate-associated RNA viruses, and invertebrate-associated RNA viruses are shown in black, purple and green, respectively. In all cases, mean values within each viral family were compared by using a one-way ANOVA, followed by Šídák’s multiple comparisons test (ns = not significant with *P* >0.05; \**P* < 0.05; \*\**P* < 0.01).

To determine whether the differences in genome size may be associated differences in the size of the gene that dictates virus replication, we compared the sizes of gene encoding the RNA-dependent RNA polymerase (RdRp) – a defining characteristic of RNA viruses – within each viral order. In some positive-stranded RNA viruses (i.e., phyla *Pisuviricota*, and *Kitrinoviricota*) the RdRp is part of the polyprotein, making an accurate comparison of the gene encoding RdRp difficult. In the *Negarnaviricota* and *Duplornaviricota*, where the start and stop codons for the RdRp gene can be clearly determined, we examined the size of the gene encoding RdRp. This analysis revealed that the size of the RdRp gene in invertebrate-associated viruses was significantly larger than in vertebrate viruses in the *Bunyavirales* (*p* < 0.0001), while there were no significant differences in the *Mononegavirales, Articulavirales, Reovirales* and *Ghabrivirales*) (Fig. 3C). To examine these differences in more detail, we compared the genome size and the gene size encoding RdRp within each viral family in the *Bunyavirales* and *Mononegavirales*. In the *Phenuiviridae* (*Bunyavirales*), invertebrate-associated RNA viruses have significantly larger RdRp genes than vertebrate viruses (*p* = 0.0183) (Fig. 3A and 3D) and slightly larger genomes than vertebrate viruses, although this difference was not significant (*P* = 0.9186). In contrast, in the *Rhabdoviridae* (*Mononegavirales*) there were no significant differences in the size of genome and the RdRp gene between invertebrate- and vertebrate-associated viruses (*p* = 0.0627, *p* = 0.0551, respectively) (Fig. 3B and 3E). However, there was a strong correlation between genome size and the size of the gene encoding RdRp in several groups (Supplementary fig. 3). In the *Bunyavirales* there was a strong correlation in both vertebrate- (R^2^ = 0.6469, *P* < 0.0001) and invertebrate-associated viruses (R^2^ = 0.7284, *P* < 0.0001) (Supplementary fig. 3), while in the *Mononegavirales* there was a strong correlation in vertebrate-associated viruses (R^2^ = 0.5946, *p* < 0.0001), although not for invertebrate-associated viruses (R^2^ = 0.1566, *p* < 0.0001) (Supplementary fig. 3). Hence, these results suggest that the increased gene size of RdRp may be one factor that contributes to genome size expansion, although this differs between virus orders and families.

We also compared the genome sizes between segmented and non-segmented viruses within each viral order. Some orders of RNA viruses contain both segmented and non-segmented viruses (i.e., *Mononegavirales, Jingchuvirales, Hepelivirales, Amarillovirales, Nodamuvirales* and *Nidovirales*), while others only contain segmented viruses (i.e., *Articulavirales, Bunyavirales, Muvirales, Goujianvirales, Reovirales* and *Durnavirales*). The *Negarnaviricota* contain two subphyla – the *Haploviricotina* (e.g. *Mononegavirales* and *Jingchuvirales*) – that primarily comprise viruses with non-segmented genomes, and the *Polyploviricotina* (e.g. *Articulavirales*, and *Bunyavirales*) that largely consist of viruses with segmented genomes (Koonin et al. 2020; Kuhn et al. 2019; Siddell et al. 2019). In addition, the *Duplornaviricota* have viral orders that contain only segmented or non-segmented RNA viruses. Notably, segmented viruses within the *Nidovirales* (of which there are three – Longidorus elongatus nidovirus, Hydra vulgaris nidovirus 2 and Crassostrea gigas nidovirus) had significantly larger genome sizes than non-segmented viruses (Fig. 1C; Supplementary table 1). Conversely, there were no significant differences in the size of the segmented and non-segmented genomes in the *Mononegavirales, Jingchuvirales, Hepelivirales, Amarillovirales* and *Nodamuvirales* (Fig. 1C). Hence, these results again suggest that whether viral genome segmentation contributes to an increase in viral genome size depends on taxonomic group.

Next, we compared genome sizes between viruses with differing numbers of segments. In the *Bunyavirales*, viruses with three or more genome segments had significantly larger genome sizes than viruses with two segments (*p* < 0.05) (Fig. 1C), while viruses with six segments had significantly larger genome sizes than viruses with three segments (*p* = 0.0039) (Fig. 1C). In the case of the *Reovirales*, Nephila clavipes virus 6, which has two genome segments, had a significantly smaller genome than viruses with eight, nine, ten, 11 or 12 segments (*p* = 0.0018, *p* < 0.0001, *p* = 0.0003, *p* = 0.0005 and *p* < 0.0001, respectively), while there was no significant difference in genome size among viruses with more than seven segments (*p* > 0.3) (Fig. 1C). In the *Articulavirales*, there were no significant differences in genome size among viruses possessing any number of genome segments (*p* > 0.9) (Fig. 1C). For the *Articulavirales* we also examined whether the average size of segments differed among viruses possessing six, seven, eight or ten genome segments. Accordingly, viruses with eight segments had significantly smaller average segment sizes than those with six segments but not those with seven segments (*P* < 0.0001 and *P* = 0.0805, respectively), while the average size of segments in Tilapia lake virus that has ten genome segments were significantly smaller than viruses that possessed fewer segments (*p* < 0.0001 in all cases) (Supplementary fig. 4). Hence, overall genome sizes are maintained despite differences in the number of genome segments.

We also examined patterns of genome segmented between vertebrate- and invertebrate-associated RNA viruses in each viral order (the exceptions were the *Jingchuvirales, Muvirales* and *Goujianvirales* that only contained invertebrate-associated viruses). In our analysis, 188 invertebrate-associated viruses contained segmented genomes compared to 184 vertebrate viruses. In the *Articulavirales*, vertebrate RNA viruses had more segments than invertebrate RNA viruses (Supplementary fig. 5). Similarly, in the *Bunyavirales* and *Reovirales*, invertebrate RNA viruses had more segments than vertebrate RNA viruses (Supplementary fig. 5). In the *Mononegavirales, Hepelivirales, Amarillovirales* and *Nidovirales*, segmented genomes were only found in invertebrate-associated RNA viruses (Supplementary fig. 5), while in the *Nodamuvirales*, non-segmented genomes were found only in invertebrate RNA viruses. Lastly, in the *Durnavirales*, all viruses had bi-segmented genomes, such that there were no differences in the number of segments between invertebrate- and vertebrate-associated RNA viruses (Supplementary fig. 5).

Finally, we analyzed the association between the genome length of segmented and non-segmented viruses, and to that of the RdRp gene, in the *Negarnaviricota* and *Duplornaviricota*. In the case of the *Negarnaviricota* there was no significant difference in genome size between viruses with segmented and non-segmented genomes (*p* = 0.2525) (Supplementary fig. 6), although the RdRp gene was significantly larger in viruses with segmented than non-segmented genomes (*p* < 0.0001). Similar results were observed even if the RdRp gene of the *Articulavirales* was assumed to represent PB1 only (*p* = 0.2512 and *p* = 0.0008 for genome size and RdRp gene size, respectively) (Supplementary fig. 7C). In contrast, in the *Duplornaviricota*, genome size was significantly larger in segmented than non-segmented genome viruses (*p* = 0.0032), while there was no significant difference in size of the RdRp gene between segmented and non-segmented viruses (*p* = 0.1859) (Supplementary fig. 6).

To assess the relative importance of individual features – host (i.e. invertebrate or/and vertebrate), viral order (18 viral orders) and genome segmentation (i.e., segmented or non-segmented) – in determining virus genome size, we used a random forest regression model to analyze the genome size data to be linked to each information. We first calculated “feature importance” which provides information on the relative significance of each covariate in building the model. This analysis revealed that viral order (particularly the *Nidovirales, Reovirales, Picornavirales, Stellavirales* and *Nodamuvirales*) was an important feature in determining genome size (Fig. 4A). In the other virus orders, the feature importance of viral order was similar to or lower than that of the genome segmentation (i.e., segmented or non-segmented) or host group (i.e. invertebrate or/and vertebrate) (Fig. 4A). To evaluate the contribution of each feature (viral order, host, and segmentation) to genome size and to assess how they affected the prediction results, we calculated the sum of the absolute mean SHapley Additive exPlanations (SHAP) values for each of these features. We then compared the mean of these sums across all data points. This revealed that viral order was significantly higher in value than both segmentation and animal host (*P* < 0.0001 for both) (Fig. 4B), indicating that it contributes most to genome size. Overall, therefore, viral order has a greater impact in determining the genome size of RNA viruses that genome segmentation or major animal host.

**Figure 4.**
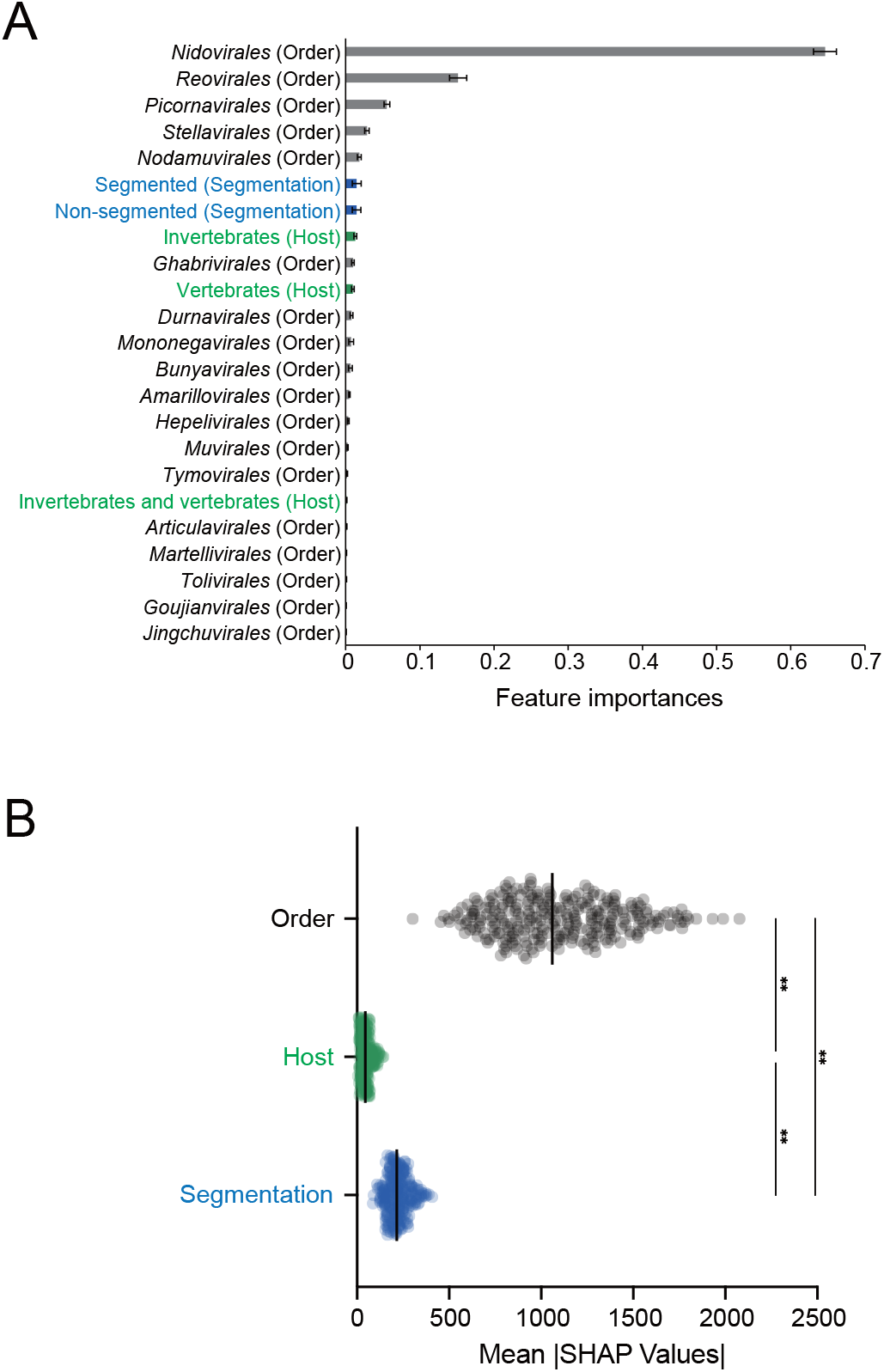
Statistical importance of viral order, host, and segmented genome in determining the genome sizes of animal RNA viruses. (A) Mean feature importance in relation to genome size. Data are shown as the mean of 100 replications. Error bars indicate the standard deviation. The features included in each qualitative trait’s groups (i.e., virus order, host, and segmentation group) are shown in gray and green and blue, respectively. Each feature is ordered from top to bottom based on its mean importance, with the highest at the top. (B) Evaluating contributions of qualitative traits to genome size using mean SHAP values. The mean of the sum of the absolute mean SHAP values for all samples (data points) was calculated for each of the qualitative traits: viral order, host, and segmentation. The mean values were compared by using a one-way ANOVA, followed by Tukey’s multiple comparisons test (**P < 0.01).

## Discussion

We performed a detailed analysis of the variation in genome sizes, and its determinants, within and among different orders and families of animal RNA viruses. Our results indicate that the genome sizes of animal RNA viruses exhibit significant variation both within and between different viral orders. In particular, the *Nidovirales*, which includes the coronaviruses, had a far greater variation in genome size than other viral orders. The larger genome sizes for at least some viruses in this group are likely explained by the presence of the ExoN domain that reduces the mutation rate and enables genomes to expand in size without accumulating excessive numbers of deleterious mutations. Indeed, it is notable that the smallest nidovirus – Equine arteritis virus at 12,704 nt – lacks an ExoN domain (Saberi et al. 2018). However, it is also of note that viral genomes of comparable size to those of the *Nidovirales* have recently been detected in the *Flaviviridae*, yet which seem to lack a detectable ExoN domain (Petrone et al. 2024). Hence, there might be other evolutionary solutions for RNA viruses to overcome the notional error threshold. In this context it is notable that the genomes of invertebrate-associated viruses were significantly larger than those associated with vertebrates in both the *Nidoviridae* and the *Amarillovirales* (i.e., *Flaviviridae*). Thus, a better understanding of the mechanisms by which large viruses overcome mutational burdens might be found in further studies of invertebrate-associated viruses.

We also found a complex relationship between RNA virus genome size and major animal host type and pattern of genome segmentation. In most cases there were no significant differences in the genome sizes of vertebrate or invertebrate viruses, although when there were differences invertebrate viruses tended to be associated with longer genomes (i.e., the *Bunyavirales, Amarillovirales, Nidovirales, Picornavirales*) with the *Mononegavirales* showing the opposite pattern. Although the reasons for the increased genomes in the vertebrate *Mononegavirales* are unclear, it is striking that this size increase was associated with multiple families (i.e., the *Filoviridae, Paramyxoviridae, Pneumoviridae* and *Sunviridae*) that only contain vertebrate viruses. In addition, it is notable that the longest invertebrate-associated picornavirus in the order *Picornavirales* (12,333 nt) did not exceed the smallest vertebrate-associated nidovirus in the order *Nidovirales* (12,704 nt), highlighting the impact of virus order. These uncertainties notwithstanding, it is striking that the viruses of vertebrates do not follow their hosts in generally being larger and more complex than those of invertebrates.

Any association between genome segmentation and genome size similarly seemed to depend on viral order. In the case of positive-stranded RNA viruses, within which genome segmentation has evolved multiple times, this trait was associated with larger genomes in the *Nidovirales*, while there was no difference among the *Amarillovirales, Hepelivirales* and *Nodamuvirales*. In the case of double-stranded RNA viruses and negative-stranded RNA viruses genome segmentation was associated with larger genomes in the *Duplornaviricota*, while there was no significant difference in genome size in the *Negarnaviricota*.

Interestingly, segmented genomes were only found in invertebrate-associated RNA viruses in the *Mononegavirales, Hepelivirales, Amarillovirales* and *Nidovirales*. It has been proposed that various aspects of virus morphology, such as the size and shape of viral particles, impact aspects of virus assembly and in so doing constrain the sizes of virus genomes. For example, while influenza A and B viruses have eight segment genomes and influenza C and D viruses have seven segment genomes, all can incorporate eight RNA segments within their particles (Nakatsu et al. 2016; Nakatsu et al. 2018; Noda et al. 2006; Noda et al. 2012). Our results show that in the *Articulavirales* (that includes influenza viruses), six segment viruses have significantly larger average segment sizes than seven and eight segment viruses, although there is no association between segment number and genome size. Hence, there may have been selection for an optimal number and size of segments within the range that can be stably packaged into particles, yet which fall within an upper limit on overall genome size. In support of this, Tilapia lake virus has a ten segment genome, but is slightly smaller in size than *Articulavirales* with eight segment genomes (although the difference is not significant) and with significantly smaller genome segments. How the segments of Tilapia lake virus are packaged is unknown, but it is possible that by reducing segment sizes, Tilapia lake virus might overcome the constraints on the number of RNA segments for stable packaging. Likewise, in the *Reovirales*, there were no significant differences in genome size among viruses containing between seven and 12 segments, suggesting that there are overall constraints on genome size irrespective of the number of genome segments. However, genome size and capsid volume are not proportional in RNA viruses compared to DNA viruses (Chaudhari et al. 2021). For example, the capsid volume of RNA viruses can vary by a factor of ∼1000, while their genome size varies by only ∼10 fold (Chaudhari et al. 2021). This suggests that the constraints imposed by genome size and size of the viral particles again depends on higher virus taxonomy. Since the morphological characteristics or mechanics of packaging of the increasingly large number of viruses discovered through metagenomic analysis are not yet well understood, it is evident that a better a biophysical characterization of newly discovered viruses is warranted.

A common theory in the evolutionary biology of cellular organisms is that there are important historical – that is, phylogenetic – constraints on adaptive flexibility which hinders the evolution of optimal adaptive solutions (Gould and Lewontin 1979). Hence, organismal evolution is contingent on the traits inherited from their ancestors, and no organism exists in a completely adaptively optimized condition (as such an organism would live forever and have infinite off-spring). These constraints can hinder evolution in other directions. For example, the number of cervical vertebrae in mammals (typically seven) is tightly constrained by genetic interactions during development (Galis et al. 2006; Narita and Kuratani 2005). As our random forest regression analysis indicated that RNA virus order has a greater impact on virus genome size than major host group or pattern of segmentation, it is possible that this theory can be extended to viruses. Hence, the effect of the host on genome size needs to be understood for each viral order which have inherited a particular set of virological traits.

An important limitation of our study is the very high number of undiscovered viruses, that will number many millions, especially in invertebrate taxa that have only been poorly sampled to date. Recently, metagenomic analyses of various invertebrates have detected viral genomes that greatly extend our current understanding of RNA virus genomes and their evolution (Neuman et al. 2024; Petrone et al. 2024). It is therefore conceivable that viral genomes sizes will continue to expand with greater sampling and challenge the observations made here. In addition, there remains controversy regarding the viral taxonomic ranks above the phyla level or order level (Simmonds et al. 2023), limiting the scope of evolutionary discussions in this study in the kingdom *Orthornavirae* in the realm *Riboviria*. The discovery of more novel viruses from diverse hosts will not only help to fill gaps in virus taxonomy, but provide a better understanding of the evolution of genome size of RNA viruses.

## Materials and Methods

### Sequence Collection

Complete genome sequences of RNA viruses were obtained from the Virus Genome Resource (https://www.ncbi.nlm.nih.gov/genomes/GenomesGroup.cgi?taxid=10239&sort=taxonomy) in May 2024. Partial sequences and those without host annotation were excluded. We supplemented these data with viral sequences collated from the National Center for Biotechnology Information (NCBI)/GenBank database and a relevant publication (Neuman et al. 2024). Accordingly, the total number of RNA viruses with complete genomes analyzed in this study was 1467, representing 18 orders (including 58 family) of RNA viruses (Supplementary table 1). The genome sizes of viruses with segmented genomes were calculated by summing the sizes of each genome segment. For analyses of the RdRp gene alone, we added 92 viruses, comprising 18 from the *Articulavirales*, 41 from the *Bunyavirales*, 12 from the *Mononegavirales* and 21 from the *Jingchuvirales* (Supplementary table 2). The length of the RdRp genes of viruses in the *Negarnaviricota* and *Duplornaviricota* were measured from the start to the stop codon (Supplementary table 1, Supplementary table 2). In the case of the *Articulavirales* which have a complex of replication proteins, two methods were used to calculate the size of the RdRp: (i) summing the sizes of three genes that encode polymerase basic protein 1 (PB1), polymerase basic protein 2 (PB2), and polymerase acidic protein (PA); and (ii) utilizing the PB1 gene only (Supplementary fig. 7).

### Statistical Analysis

Values of skewness and kurtosis were computed to assess the asymmetry and sharpness of the distribution. To calculate skewness and kurtosis for the sample size *n*, we subtracted the mean value 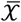 from each data 𝒳_*i*_ and standardized the result by dividing by the standard deviation *s*. For small samples, bias corrections were applied.

Skewness is adjusted using a correction factor 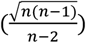:

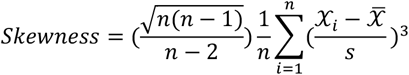

Kurtosis is calculated as:

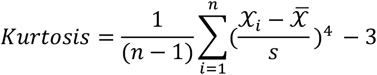

A homogeneity of variance test was performed by using mean-based Levene’s test (Draper and Hunter 1969), which is the one-way analysis of variance F-test on 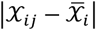, the absolute deviations of the 𝒳_*ij*_ from their group mean 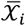. Mean values were compared by using a one-way ANOVA, followed by Tukey’s multiple comparisons test. A *p*-value < 0.05 denoted statistical significance.

We used linear regression to determine whether there was a correlation between viral genome size and the size of the gene encoding the RdRp. Accordingly, linear regression graphs were obtained using Graph Prism 10 software (version 10.3.1). The statistical significance of each variable was evaluated with a *p*-value (*p* < 0.05 was considered significant), and the goodness of fit of the overall model was indicated by R^2^. To compare the genome size or size of the gene encoding the RdRp, the mean values within each viral order were compared by using a one-way ANOVA, followed by Šídák’s multiple comparisons test, with *p*-value < 0.05 again denoting statistical significance. All statistical analysis were performed using Graph Prism 10 software (version 10.3.1) and in-house python scripts.

### Random Forest Regression Model

A Random Forest Regression was employed to model the relationship between the features and the target variable, in this case genome size. Various categorical features, including host (i.e. invertebrate or/and vertebrate), viral order (18 viral orders) and genome segmentation (i.e., segmented or non-segmented), were transformed into dummy variables using the “pd.get_dummies” function. The features and target variable were then separated, and feature scaling was performed using the StandardScaler to ensure all features were on a similar scale. The Random Forest model was performed using the scikit-learn library (Lars Buitinck 2013) in Python (version 3.10.14). The data was randomly split into training (80%) and testing (20%) sets for each iteration of the model training, which was repeated 100 times. During each iteration, feature importances were recorded, and five-fold cross-validation was conducted to evaluate the model’s performance, utilizing mean squared error (MSE) as the scoring metric. This approach involved dividing the training data into five subsets, training the model on four subsets, and validating it on the remaining subset. This process was repeated five times, with each subset serving as the validation set once, resulting in an average MSE across all folds. Additionally, SHapley Additive exPlanations (SHAP) values (Lundberg 2017) were computed using SHAP TreeExplainer library to quantify the impact of each feature on the target variable. The results, including the mean and standard deviation of feature importances and SHAP values, were compiled into a Pandas DataFrame and saved in an Excel file for further analysis.

## Acknowledgments

We thank Dr. Chris Lauber for sharing genome information on the viruses in the order *Nidovirales*. We also thank Ms. Kanako Hiromatsu and Dr. Tokiko Watanabe for their support with grant and equipment procedures at Osaka University. This study was supported by RIKAKEN HOLDINGS CO. Young Researcher Support Grant-in-aid (to K.T.).

## Author contributions

K.T. and E.C.H. conceived the study. K.T. analyzed data. K.T. and E. C.H. wrote the manuscript and approved the final manuscript.

## Data availability

The data sets used in this study are provided in the Supplementary tables 1 and 2.

## Supplementary information

**Supplementary Figure 1. Comparison of genome size variation in each viral order using Levene’s Test**. The mean of the absolute deviation of each virus calculated from the mean in each viral order is shown. Mean values were compared using a one-way ANOVA, followed by Tukey’s multiple comparisons test (\**P* < 0.05; \*\**P* < 0.01; \*\*\**P* < 0.001; \*\*\*\**P* < 0.0001)

**Supplementary Figure 2. Genome size of animal RNA viruses within each viral family within the orders *Amarillovirales, Nidovirales* and *Picornavirales***. (A-C) The genome sizes of RNA viruses in each viral family belonging to the *Amarillovirales*, (A), *Nidovirales* (B) and *Picornavirales* (C). Vertebrate-associated RNA viruses, both vertebrate- and invertebrate-associated RNA viruses, and invertebrate-associated RNA viruses are shown in black, purple and green, respectively. The mean values within each viral order were compared by using a one-way ANOVA, followed by Šídák’s multiple comparisons test (ns = not significant with *P* >0.05; \*\**P* < 0.01, \*\*\*\**P* < 0.0001).

**Supplementary Figure 3. Correlations between genome size and the size of the gene encoding the RdRp**. Correlations between genome size and gene size encoding RdRp in each viral order. Vertebrate-RNA viruses and invertebrate-associated RNA viruses are shown in black and green, respectively. Simple linear regressions of each group were shown.

**Supplementary Figure 4. Comparison of genome segment size in RNA viruses with six, seven, eight or ten segments from the order *Articulavirales***. The average size of genome segments in viruses with six, seven, eight or ten segments within the *Articulavirales* is shown. Data are shown as the mean size of all segments in each virus group, with the average value for each segment indicated by a dot. Error bars indicate the standard deviation. The mean values of all segments in each virus group were compared by using a one-way ANOVA, followed by Tukey’s multiple comparisons test (ns = not significant with *P* >0.05; \*\**P* < 0.01; \*\*\*\**P* < 0.0001).

**Supplementary Figure 5. Number of segments in vertebrate- and invertebrate-associated RNA viruses in each viral order**. Vertebrate-associated RNA viruses, both vertebrate- and invertebrate-associated RNA viruses, and invertebrate-associated RNA viruses are shown in black, purple and green, respectively.

**Supplementary Figure 6. Genome sizes and the gene size of the RdRp in segmented and non-segmented RNA viruses of the phyla *Negarnaviricota* and *Duplornaviricota***. The genome size and RdRp gene size of non-segmented and segmented RNA viruses belonging to each phylum. Non-segmented and segmented RNA viruses are shown in black and blue, respectively. The mean values within each viral order were compared by using a Kruskal-Wallis test, followed by Dunn’s multiple comparisons test (ns = not significant with *P* >0.05; \*\**P* < 0.01; \*\*\*\**P* < 0.0001)

**Supplementary Figure 7. Comparison of gene sizes in animal RNA viruses when the only PB1 gene of *Articulavirales* is used as the RdRp gene**. (A) Size of the gene encoding the RdRp of RNA viruses in each viral order belonging to the *Negarnaviricota* and *Duplornaviricota*. Vertebrate-associated RNA viruses, RNA viruses associated with both vertebrates and invertebrates, and invertebrate-associated RNA viruses are shown in black, purple and green, respectively. The mean values within each viral order were compared by using a one-way ANOVA, followed by Šídák’s multiple comparisons test (ns = not significant with *P* >0.05; \*\*\**P* < 0.001; \*\*\*\**P* < 0.0001). (B) Correlations between genome size and gene size encoding RdRp in the *Articulavirales*. Vertebrate- and invertebrate-associated RNA viruses are shown in black and green, respectively. Note: RNA viruses associated with both vertebrates and invertebrates were included in the vertebrate RNA viruses. Simple linear regressions of each group were shown. (C) Genome size and RdRp gene size of non-segmented and segmented RNA viruses belonging to the *Negarnaviricota*. Non-segmented and segmented RNA viruses are shown in black and blue, respectively. The mean values within each viral order were compared by using a Kruskal-Wallis test, followed by Dunn’s multiple comparisons test (ns = not significant with *P* >0.05; \*\**P* < 0.01; \*\*\*\**P* < 0.0001)

**Supplementary Table 1. List of viruses used for comparisons of genome size in RNA viruses**.

**Supplementary Table 2. List of viruses added for the comparison of RdRp gene sizes**.

## References

Aparicio, S., et al. (2002) ‘Whole-Genome Shotgun Assembly and Analysis of the Genome of Fugu rubripes’, Science, 297: 1301–10.

Biscotti, M. A., Olmo, E., and Heslop-Harrison, J. S. (2015), ‘Repetitive DNA in Eukaryotic Genomes’, Chromosome Research, 23: 415–20.

Boehm, T. (2012) ‘Evolution of Vertebrate Immunity’, Current Biology, 22, R722–32.

Canapa, A., et al. (2015) ‘Transposons, Genome Size, and Evolutionary Insights in Animals’, Cytogenetic and Genome Research, 147: 217–39.

Chao, L. (1990) ‘Fitness of RNA Virus Decreased by Muller’s Ratchet’, Nature, 348, 454–5.

Chaudhari, H. V., Inamdar, M. M., and Kondabagil, K. (2021) ‘Scaling Relation Between Genome Length and Particle Size of Viruses Provides Insights into Viral Life History’, iScience, 24: 102452.

Davidson, E. H. and Erwin, D. H. (2009) ‘An Integrated View of Precambrian Eumetazoan Evolution’, Cold Spring Harbor Symposia on Quantitative Biology, 74: 65–80.

dos Reis, M., et al. (2015) ‘Uncertainty in the Timing of Origin of Animals and the Limits of Precision in Molecular Timescales’, Current Biology, 25: 2939–50.

Draper, N. R., and Hunter, W. G. (1969) ‘Transformations: Some Examples Revisited’, Technometrics, 11: 23–40.

Du Pasquier, L. (2001) ‘The Immune System of Invertebrates and Vertebrates’, Comp Comparative Biochemistry and Physiology Part B: Biochemistry and Molecular Biology, 129: 1–15.

Duffy, S., Shackelton, L. A., and Holmes, E. C. (2008) ‘Rates of Evolutionary Change in Viruses: Patterns and Feterminants’, Nature Reviews Genetics, 9: 267–76.

Eckerle, L. D., et al. (2010) ‘Infidelity of SARS-CoV Nsp14-exonuclease Mutant Virus Replication is Revealed by Complete Genome Sequencing’, PLoS Pathogens, 6: e1000896.

Flajnik, M. F. and Kasahara, M. (2010) ‘Origin and Evolution of the Adaptive Immune System: Genetic Events and Selective Pressures’, Nature Reviews Genetics, 11: 47–59.

Galis, Frietson, et al. (2006) ‘Extreme Selection in Humans against Homeotic Transformations of Cervical Vertebrae’, Evolution, 60: 2643–54.

Gorbalenya, A. E., et al. (2006) ‘Nidovirales: Evolving the Largest RNA Virus Genome’, Virus Research, 117: 17–37.

Gould, S. J. and Lewontin, R. C. (1979) ‘The Spandrels of San Marco and the Panglossian Paradigm: a Critique of the Adaptationist Programme’, Proceedings of the Royal Society of Lond Series B Biological Sciences, 205: 581–98.

Gregory, T. R., (2005) ‘Genome Size Evolution in Animals’, In The Evolution of the Genome (Ed. Gregory, T. R.) pages 3–87. Academic Press.

Gregory, T.R. (2024) ‘Animal Genome Size Database’. https://www.genomesize.com/

Harvey, E. and Holmes, E. C. (2022) ‘Diversity and Evolution of the Animal Virome’, Nature Reviews Microbiology, 20: 321–34.

Holmes, E. C. (2003) ‘Error Thresholds and the Constraints to RNA Virus Evolution’, Trends in Microbiology, 11: 543–46.

Holmes, E. C. (2009) ‘The Evolution and Emergence of RNA viruses’, Oxford Series in Ecology and Evolution (OSEE). Oxford University Press, Oxford.

Kidwell, M. G. (2002) ‘Transposable Elements and the Evolution of Genome size in Eukaryotes’, Genetica, 115: 49–63.

Koonin, E. V., et al. (2020) ‘Global Organization and Proposed Megataxonomy of the Virus World’, Microbiology and Molecular Biology Reviews, 84: e00061–19.

Kuhn, J. H., et al. (2019) ‘Classify Viruses - the Gain is Worth the Pain’, Nature, 566: 318–20.

Buitinck, L., et al. (2013) ‘API Design for Machine Learning Software: Experiences from the Scikit-Learn Project’, European Conference on Machine Learning and Principles and Practices of Knowledge Discovery in Databases.

Liao, Y., et al. (2023) ‘Classification, Replication, and Transcription of Nidovirales’, Frontiers in Microbiology, 14: 1291761.

Lundberg, S. (2017) ‘A Unified Approach to Interpreting Model Predictions’, NIPS’17: Proceedings of the 31st International Conference on Neural Information Processing Systems. Pages 4768 – 77.

Nakatsu, S., et al. (2016) ‘Complete and Incomplete Genome Packaging of Influenza A and B Viruses’, mBio, 7: e01248–16.

Nakatsu, S., et al. (2018) ‘Influenza C and D Viruses Package Eight Organized Ribonucleoprotein Complexes’, Journal of Virology, 92: e02084–17.

Narita, Y. and Kuratani, S. (2005) ‘Evolution of the Vertebral Formulae in Mammals: a Perspective on Developmental Constraints’, Journal of Experimental Zoology Part B: Molecular and Developmental Evolution, 304: 91–106.

Neuman, B. W., et al. (2024) ‘RNA Genome Expansion up to 64 kb in Nidoviruses is Host Constrained and Associated with New Modes of Replicase Expression’, bioRxiv. https://www.biorxiv.org/content/10.1101/2024.07.07.602380v1.

Noda, T., et al. (2006) ‘Architecture of Ribonucleoprotein Complexes in Influenza A Virus particles’, Nature, 439: 490–2.

Noda, T., et al. (2012) ‘Three-Dimensional Analysis of Ribonucleoprotein Complexes in Influenza A Virus’, Nature Communications, 3:639.

Petrone, M. E., et al. (2024) ‘A ∼40-kb Flavi-Like Virus Does Not Encode a Known Error-Correcting Mechanism’, Proceedings of the National Academy of Sciences USA, 121:e2403805121.

Philippe, N., et al. (2013) ‘Pandoraviruses: Amoeba Viruses with Genomes Up to 2.5 Mb Reaching That of Parasitic Eukaryotes’, Science, 341: 281–86.

Pressing, J. and Reanney, D. C. (1984) ‘Divided Genomes and Intrinsic Noise’, Journal of Molecular Evolution, 20: 135–46.

Ros-Rocher, N., et al. (2021) ‘The Origin of Animals: an Ancestral Reconstruction of the Unicellular-to-Multicellular Transition’, Open Biology, 11: 200359.

Saberi, A., et al. (2018) ‘A Planarian Nidovirus Expands the Limits of RNA Genome Zize’, PLoS Pathog, 14: e1007314.

Sanjuan, R., et al. (2010) ‘Viral Mutation Rates’, Journal of Virology, 84: 9733–48.

Siddell, S. G., et al. (2019) ‘Additional changes to taxonomy ratified in a special vote by the International Committee on Taxonomy of Viruses (October 2018)’, Archives of Virology, 164: 943–46.

Simmonds, P., et al. (2023) ‘Four Principles to Establish a Universal Virus Taxonomy’, PLoS Biol, 21: e3001922.

Venkatesh, B., Gilligan, P., and Brenner, S. (2000) ‘Fugu: a Compact Vertebrate Reference Genome’, FEBS Letters, 476: 3–7.

