## Supplemental Figures for "Genome sizes of animal RNA viruses reflect phylogenetic constraints"

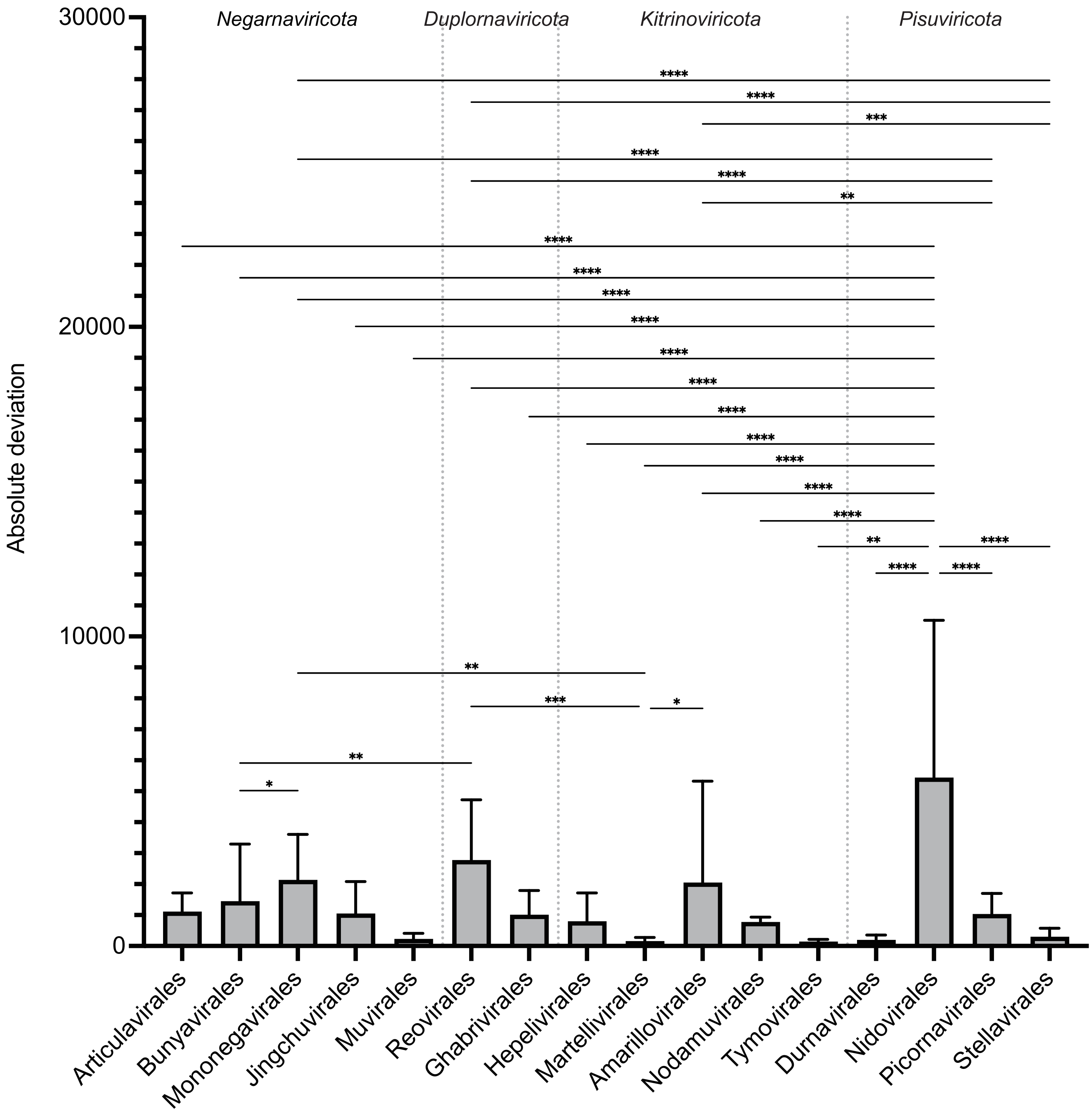


Supplementary Figure 1

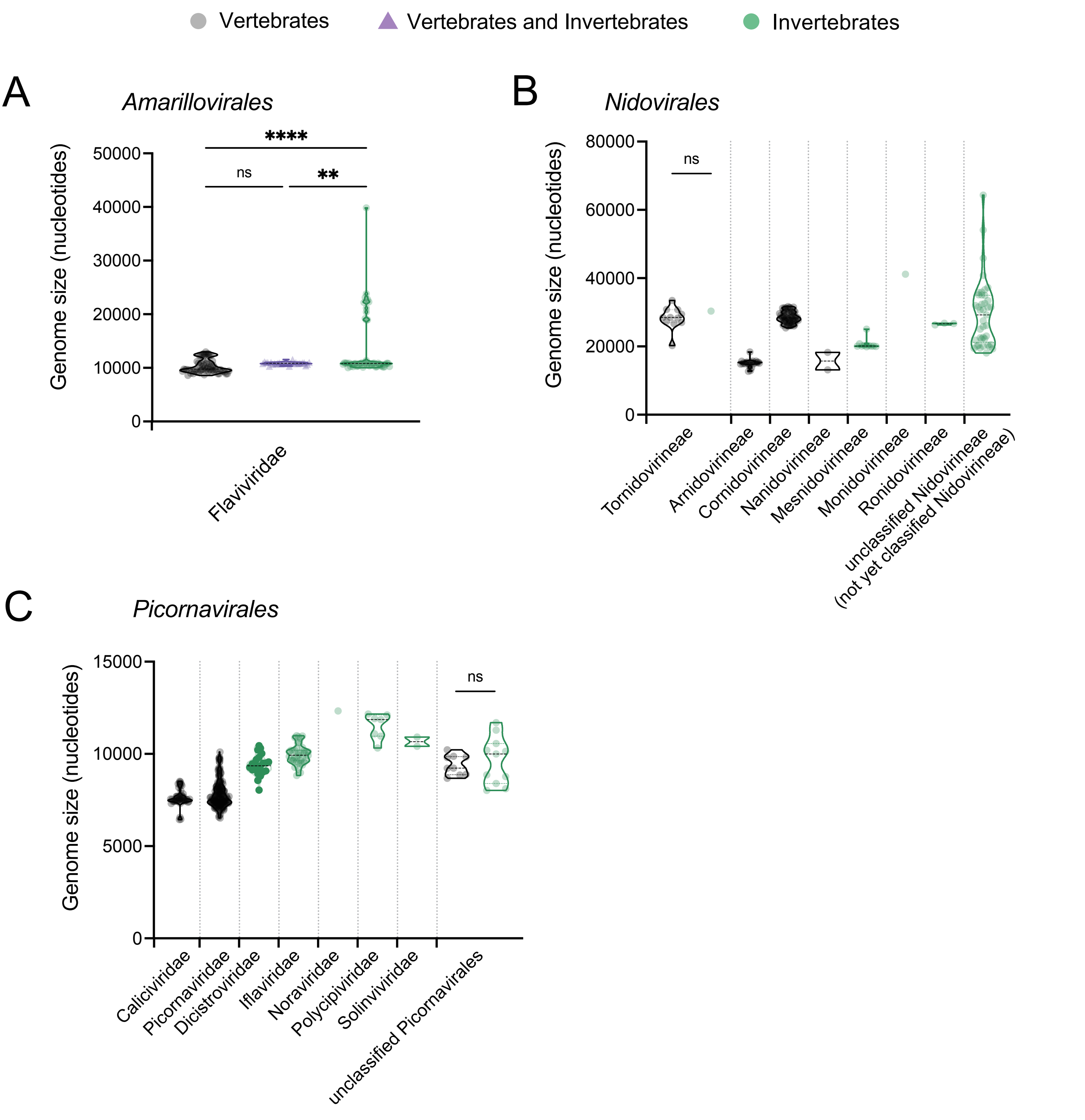


Supplementary Figure 2

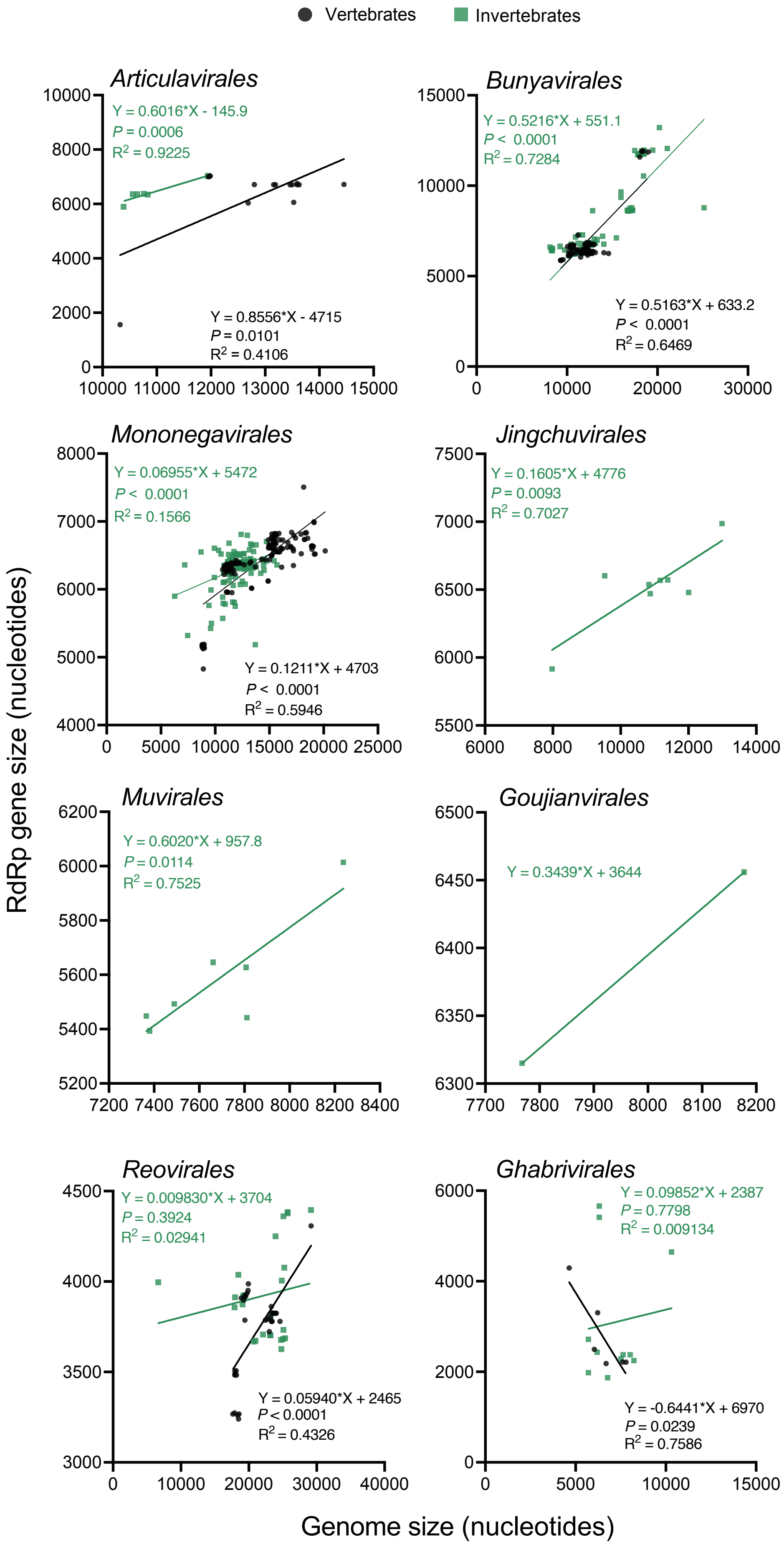


Supplementary Figure 3

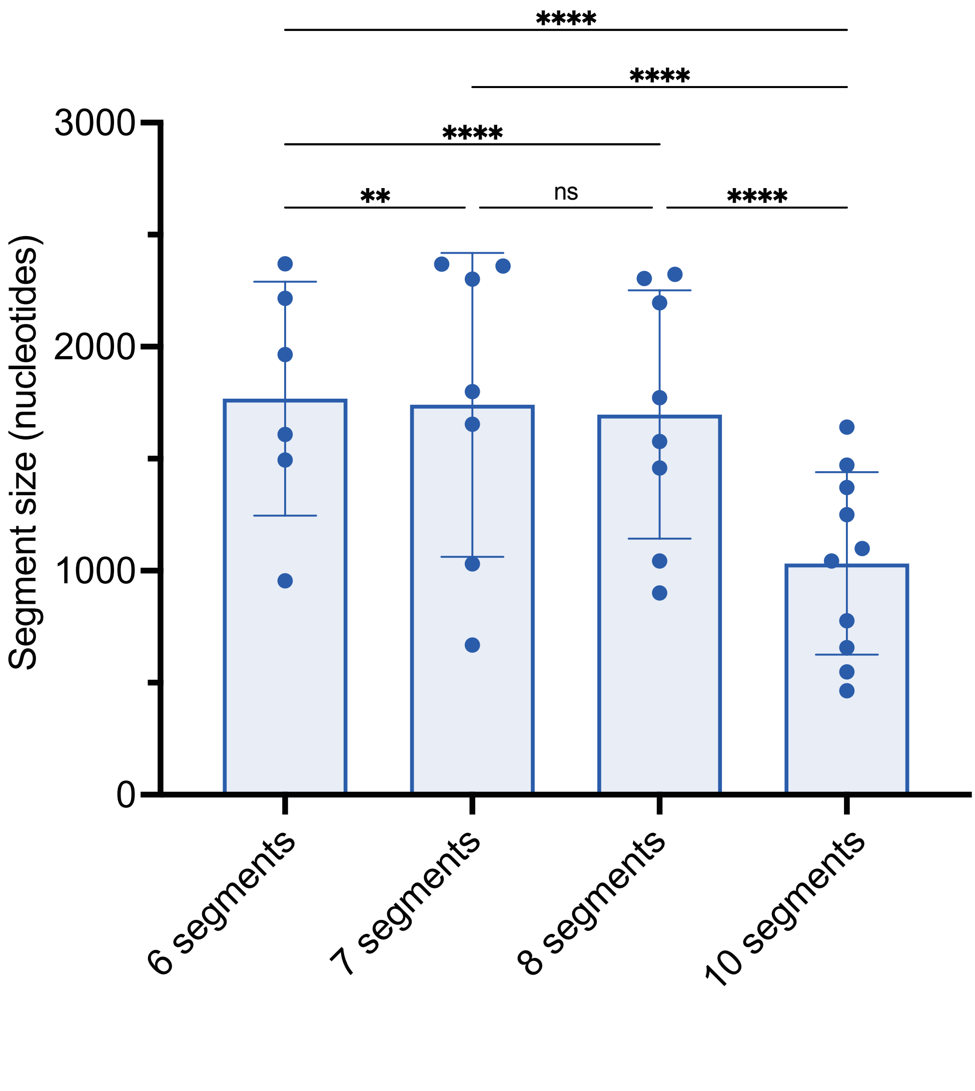


Supplementary Figure 4

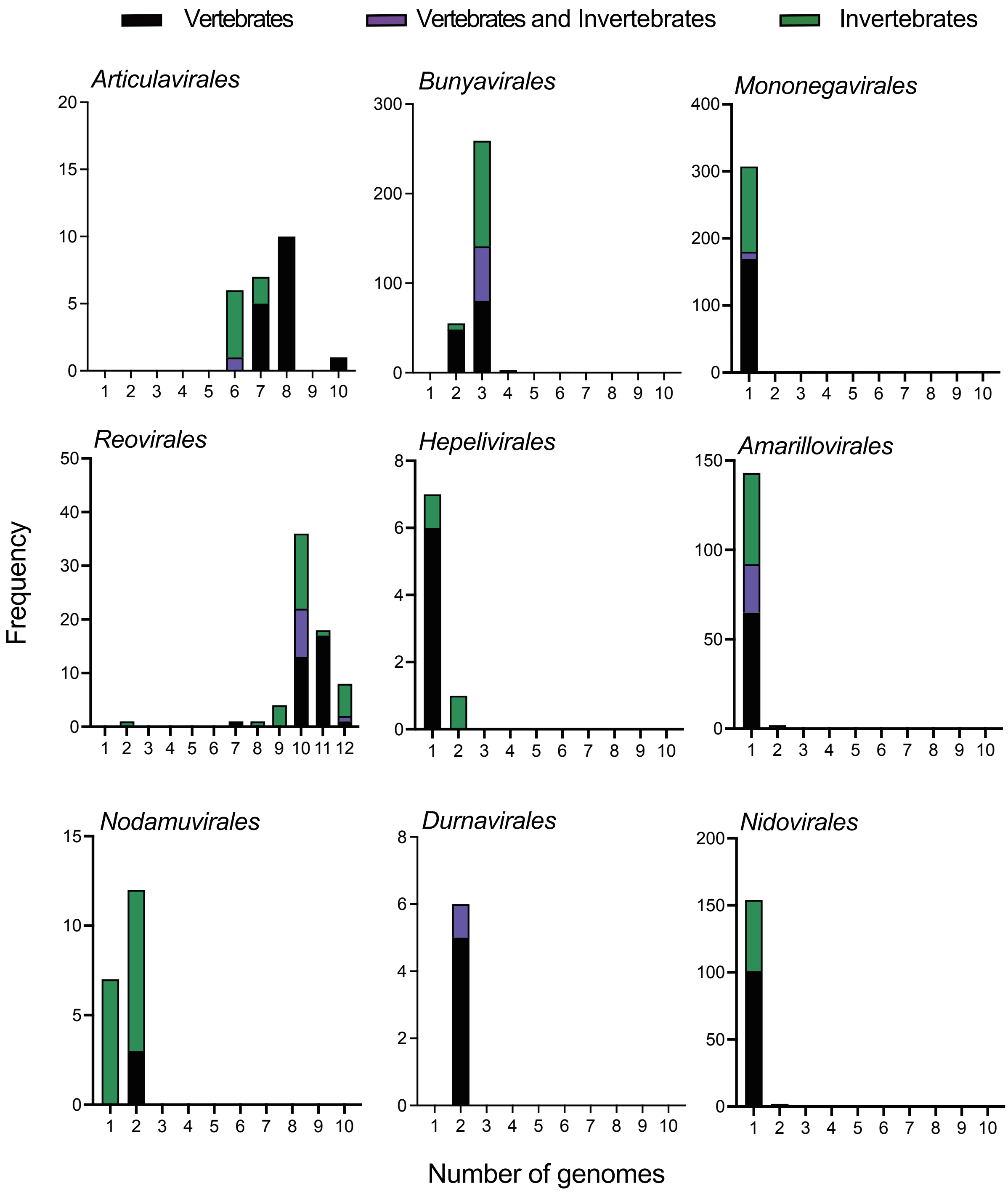


Supplementary Figure 5

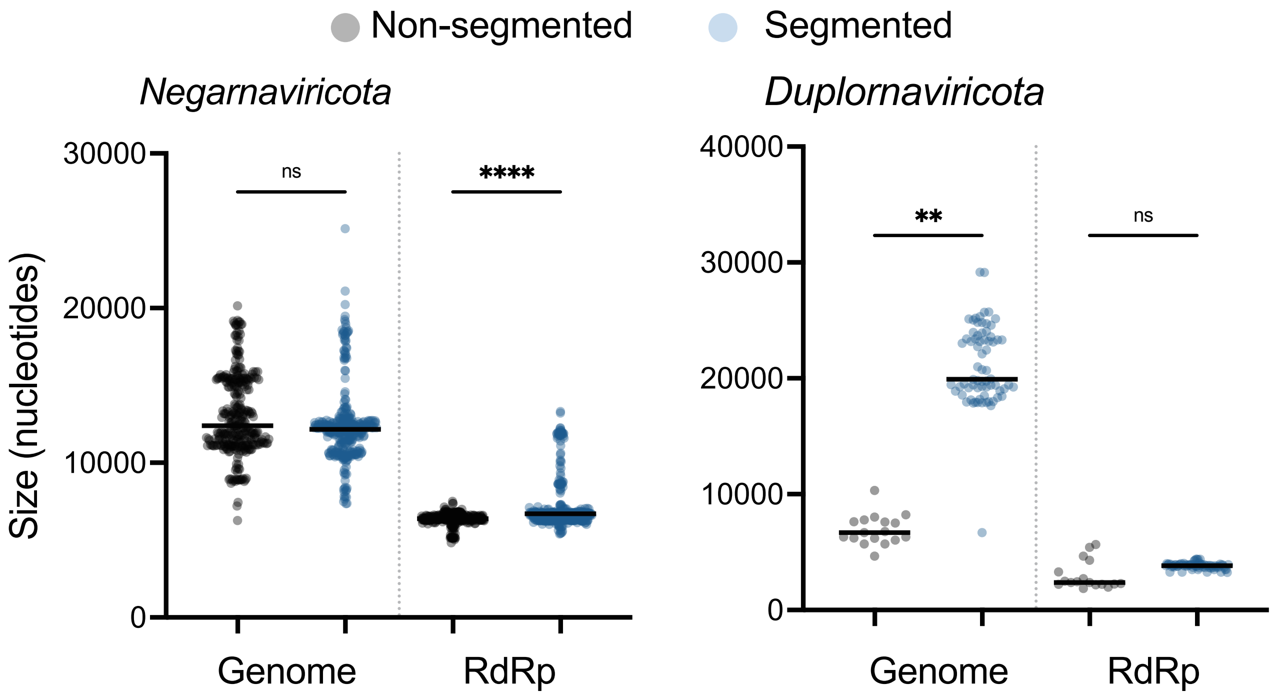


Supplementary Figure 6

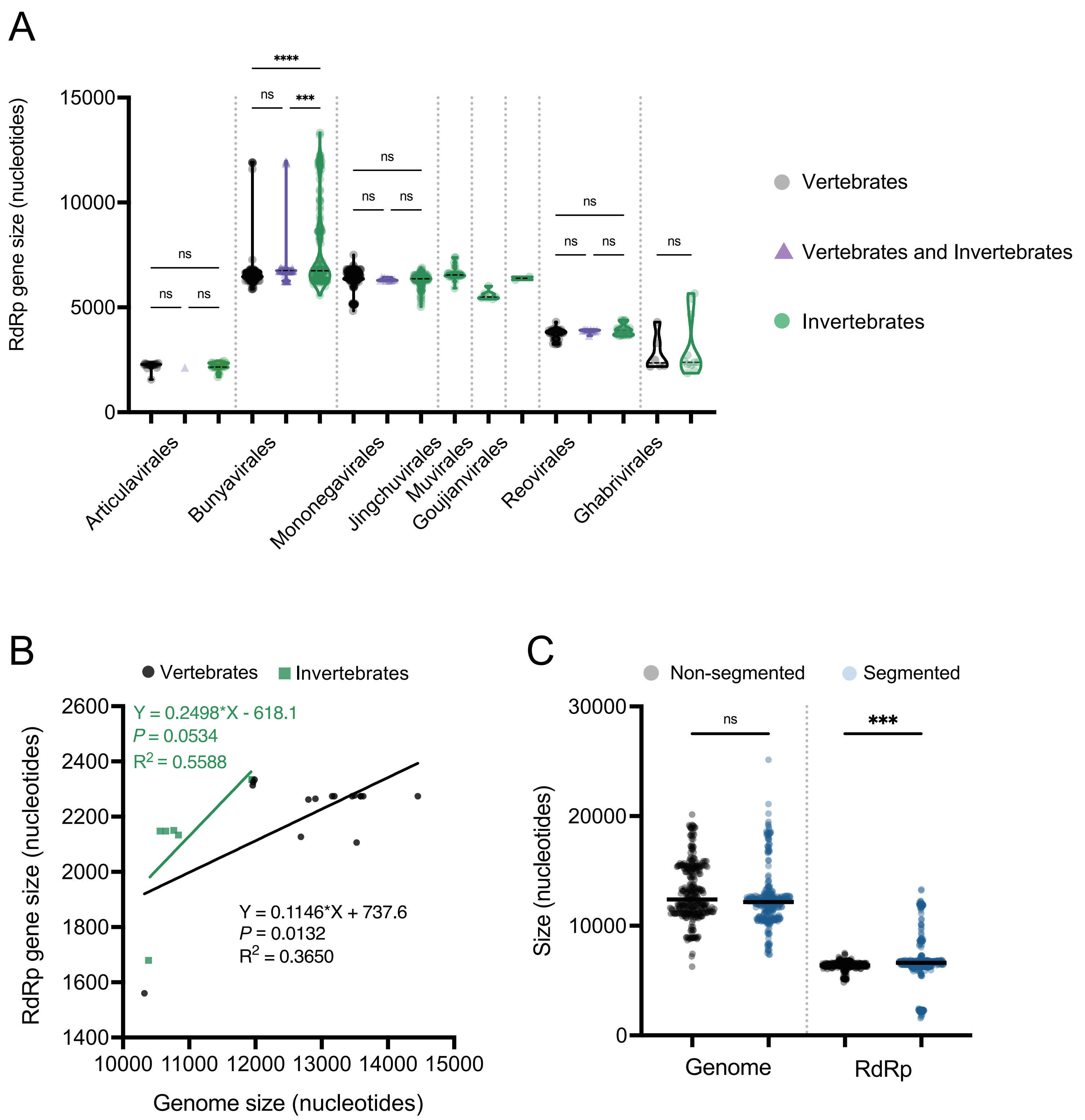


Supplementary Figure 7
